# SMARCB1 loss creates patient-specific *MYC* topologies that drive malignant rhabdoid tumor growth

**DOI:** 10.1101/2022.11.21.516939

**Authors:** Ning Qing Liu, Irene Paassen, Lars Custers, Hans Teunissen, Dilara Ayyildiz, Jiayou He, Eelco W. Hoving, Elzo de Wit, Jarno Drost

## Abstract

Malignant rhabdoid tumor (MRT) is a highly malignant and often lethal childhood cancer. MRTs are genetically defined by bi-allelic inactivating mutations in *SMARCB1*, a member of the BRG1/BRM-associated factors (BAF) chromatin remodeling complex. Mutations in BAF complex members are common in human cancer, yet their contribution to tumorigenesis remains in many cases poorly understood. Here, we studied derailed regulatory landscapes as a consequence of *SMARCB1* loss in the context of MRT. Our multi-omics approach on patient-derived MRT organoids revealed a dramatic reshaping of the regulatory landscape upon *SMARCB1* reconstitution. Chromosome conformation capture experiments subsequently revealed patient-specific looping of distal enhancer regions with the promoter of the *MYC* oncogene. This intertumoral heterogeneity in *MYC* enhancer utilization is also present in patient MRT tissues as shown by combined single-cell RNA-seq and ATAC-seq. We show that loss of *SMARCB1* drives patient-specific epigenetic reprogramming underlying MRT tumorigenesis.

## Introduction

The BRG1/BRM-associated factors (BAF) chromatin remodeling complex (or mammalian SWItch/Sucrose Non-Fermentable (SWI/SNF) complex) is a multiprotein complex with a crucial role in the regulation of transcription via nucleosome positioning. Depending on their protein composition, three main BAF complexes have been identified. The canonical BAF (cBAF) and polybromo-associated BAF (pBAF) complexes are defined by, amongst others, the SMARCB1 subunit, which is lacking in the non-canonical (nc)BAF complex^1^. Instead, the ncBAF complex contains the BRD9 and GLTSCR1/L subunits. Genes encoding members of the BAF complex are among the most commonly mutated genes in cancer, occurring in approximately 20-25% of cases^2–4^. Adult malignancies typically harbor hundreds to thousands of mutations, which complicates studying the contribution of mutations in BAF members to tumorigenesis. However, several pediatric malignancies are uniquely defined by mutations in BAF complex members^5^. Amongst those are malignant rhabdoid tumors (MRT), aggressive childhood malignancies that predominantly affect infants. They can occur throughout the body, but most commonly arise in the brain (atypical teratoid/rhabdoid tumors (AT/RT)) and kidney^6^. Despite their aggressiveness, MRTs are genetically stable tumors with a low mutational burden. In fact, in the vast majority of MRT cases (95%), bi-allelic inactivation of the BAF complex member *SMARCB1* is the only recurrent protein coding gene alteration, showing that loss of *SMARCB1* is a crucial driver event in MRT development^6,7^. We recently found that bi-allelic *SMARCB1* inactivating mutations in MRT can be shared with adjacent morphologically normal Schwann cells, suggesting that loss of *SMARCB1* is required but not sufficient for MRT development^8^.

Recent studies have demonstrated that epigenetic reprogramming, which can be caused by mutations in chromatin regulators such as BAF complex members, underpins tumor initiation and progression^9–13^. To identify and functionally study patient-specific epigenomic changes underlying malignant growth, personalized pre-clinical cell models are indispensable. Such models should reflect the epigenetic heterogeneity found between patient tumors. We and others have previously demonstrated that organoids maintain many features of the tissues they were derived from, including their epigenetic profiles, over extended serial passaging^14–17^.

Here, we apply a multi-omics approach to patient-derived organoids (PDOs) to define how loss of *SMARCB1* reorganizes chromatin and underpins MRT growth. We find patient-specific enhancer landscapes that are a consequence of a disruption in the balance of BAF complexes. Specifically, we identified patient-specific putative enhancers in different PDOs that likely regulate the expression of the *MYC* oncogene driving MRT growth. One of these putative enhancers has not been previously described within the context of tumorigenesis. The patient-specific derailed activity of these putative enhancers was also observed in patient tumors. Our study provides the first evidence that intertumoral heterogeneity in enhancer utilization drives oncogene expression and tumorigenesis in MRT.

## Results

### Analysis of chromatin dynamics in MRT PDOs reveals SMARCB1-dependent enhancer regulation

We previously showed that *SMARCB1* loss is required but not sufficient for malignant transformation^8^. We therefore hypothesized that additional epigenetic drivers are required for tumor formation. Reconstitution of the principal genetic driver event of MRT, *SMARCB1* loss, reverts MRT cells to a normalized state^8^, suggesting that epigenetic changes that have contributed to malignancy can be overcome. To find these tumor-driving regulatory changes, we lentivirally transduced a MRT PDO model (named P103)^17^ with either a control or a *SMARCB1* expression (*SMARCB1*+) plasmid and measured chromatin accessibility by assay for transposase-accessible chromatin using sequencing (ATAC-seq; Fig. 1A,B). Following *SMARCB1* reconstitution, we find 7,941 newly formed open chromatin regions (OCRs) that are enriched for transcription factor motifs from different families such as SMARCC1/2 and AP-1^18^ (Extended Data Fig. 1A,B). When we perform functional annotation of these OCRs using GREAT, we find that several categories are enriched, mostly related to differentiation and developmental processes (Extended Data Fig. 1C). For the 1,211 OCRs that were lost, the only motif that we found to be enriched was for the insulator protein CTCF (Extended Data Fig. 1B), consistent with previous reports in human embryonic stem cells and mouse embryonic fibroblasts (PMID: 31033435; PMID: 28262751).

**Fig. 1:**
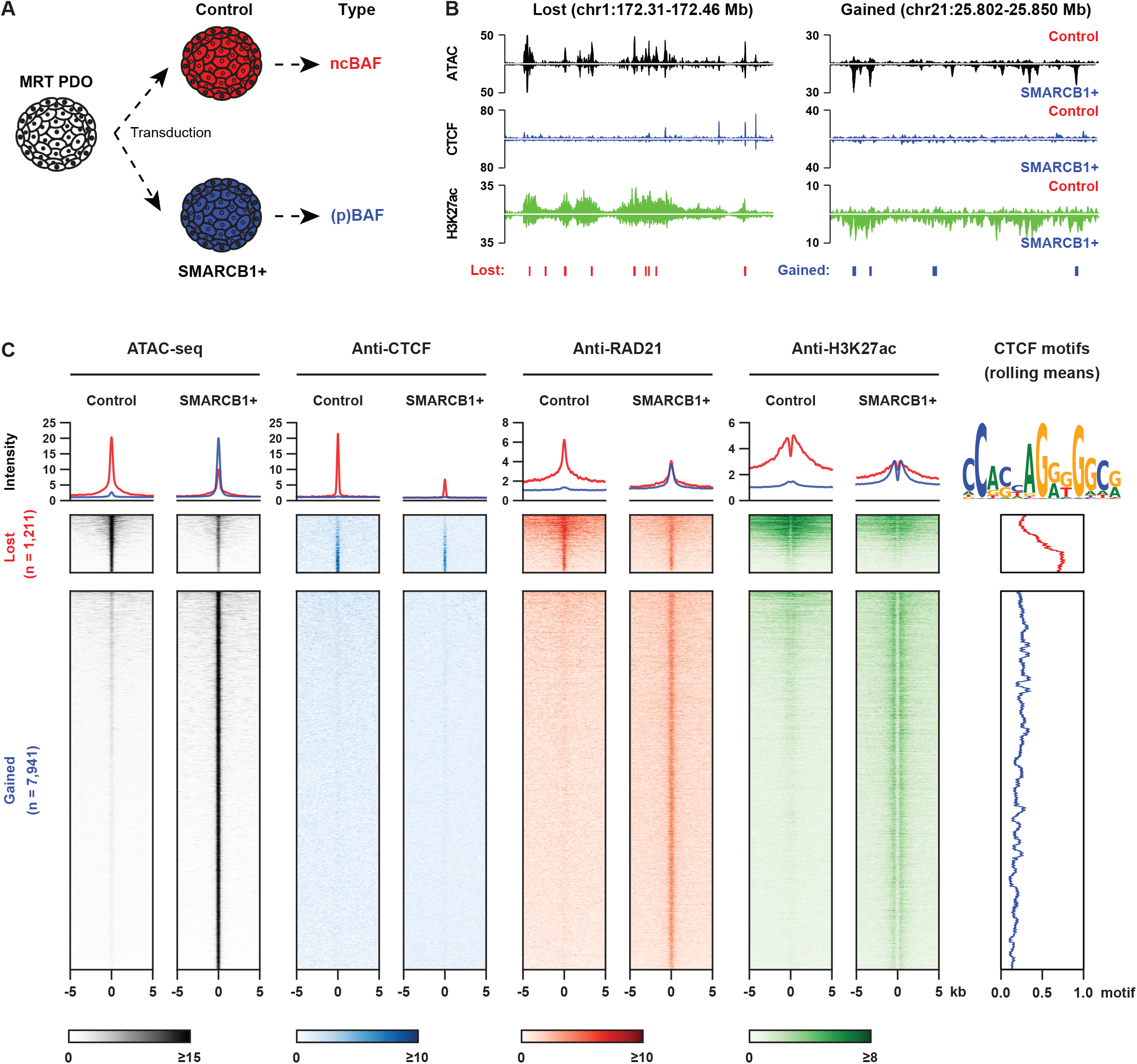
*SMARCB1* reconstitution reshapes open chromatin landscape in the MRT PDO model. (A) A MRT PDO model was transduced by a luciferase viral plasmid (control) and a viral plasmid containing the full length wild-type *SMARCB1* gene, respectively; (B) Two example loci lost (left) and gained (right) chromatin accessibility after *SMARCB1* reconstitution, respectively (ATAC-seq: n = 3, an average of the 3 replicates; ChIP-seq: n = 1); (C) Tornado-plot analysis shows that the lost (control-specific) open chromatin regions are enriched for strong enhancers and CTCF binding sites, while the gained (SMARCB1+-specific) open chromatin regions are enriched for enhancers.

ChIP-seq of CTCF showed that *SMARCB1* reconstitution results in a decrease in CTCF binding at OCRs that are lost in *SMARCB1*+ cells (Fig. 1C). In the genome, CTCF binding frequently overlaps with binding of the ring-shaped cohesin complex^19^. ChIP-seq of the cohesin complex subunit RAD21 confirmed that, in addition to decreased CTCF binding, OCRs lost in *SMARCB1+* cells show a reduction in binding of the cohesin complex (Fig. 1C). On the other hand, OCRs gained in the *SMARCB1*+ PDOs showed occupancy of RAD21 (Fig. 1C), but not CTCF. To determine how the enhancer landscape changes upon *SMARCB1* reconstitution, we performed ChIP-seq of the active enhancer mark H3K27ac. As expected, the lost OCRs show a decrease in H3K27ac levels, while a marked increase in H3K27ac levels was detected at sites that gained accessibility (Fig. 1C). Increases in enhancer marks coincides with increased binding of the cohesin complex (Fig. 1C), which is consistent with a subset of cohesin molecules binding to (super) enhancer regions, besides CTCF binding sites^20,21^. These results indicate that the canonical BAF complexes play an important role in suppressing CTCF binding to chromatin and that loss of *SMARCB1* leads to increased accessibility at CTCF-bound chromatin.

### SMARCB1 reorganizes the chromatin landscape surrounding the *MYC* oncogene

Observing a change in the CTCF and RAD21 binding landscape following *SMARCB1* reconstitution led us to hypothesize that long-distance gene regulation may be affected in these cells. To determine whether any SMARCB1-dependent changes occur in the organization of the genome (3D genome), we performed high-resolution *in-situ* Hi-C experiments^22^ in control and *SMARCB1+* P103 MRT PDOs. Despite the differences in CTCF binding to chromatin, we find that, in general, chromatin architecture is largely unaffected (Extended Data Fig. 2A,B). 3D genome features such as topologically associated domains (TADs) and A/B compartments showed limited changes upon *SMARCB1* reconstitution (Extended Data Fig. 2C-G). However, we identified 131 and 1,164 loops specific to control and *SMARCB1*+ PDOs, respectively, with a minimal 1.5-fold change of contact frequency (Extended Data Fig. 3A-D). Ranking the most prominently lost loci upon *SMARCB1* reconstitution, we found an interaction between the *MYC* oncogene and an ∼1.1 Mb distal region marked by high H3K27ac levels indicative of a super enhancer (Fig. 2A,B and Extended Data Fig. 3B).

**Fig. 2:**
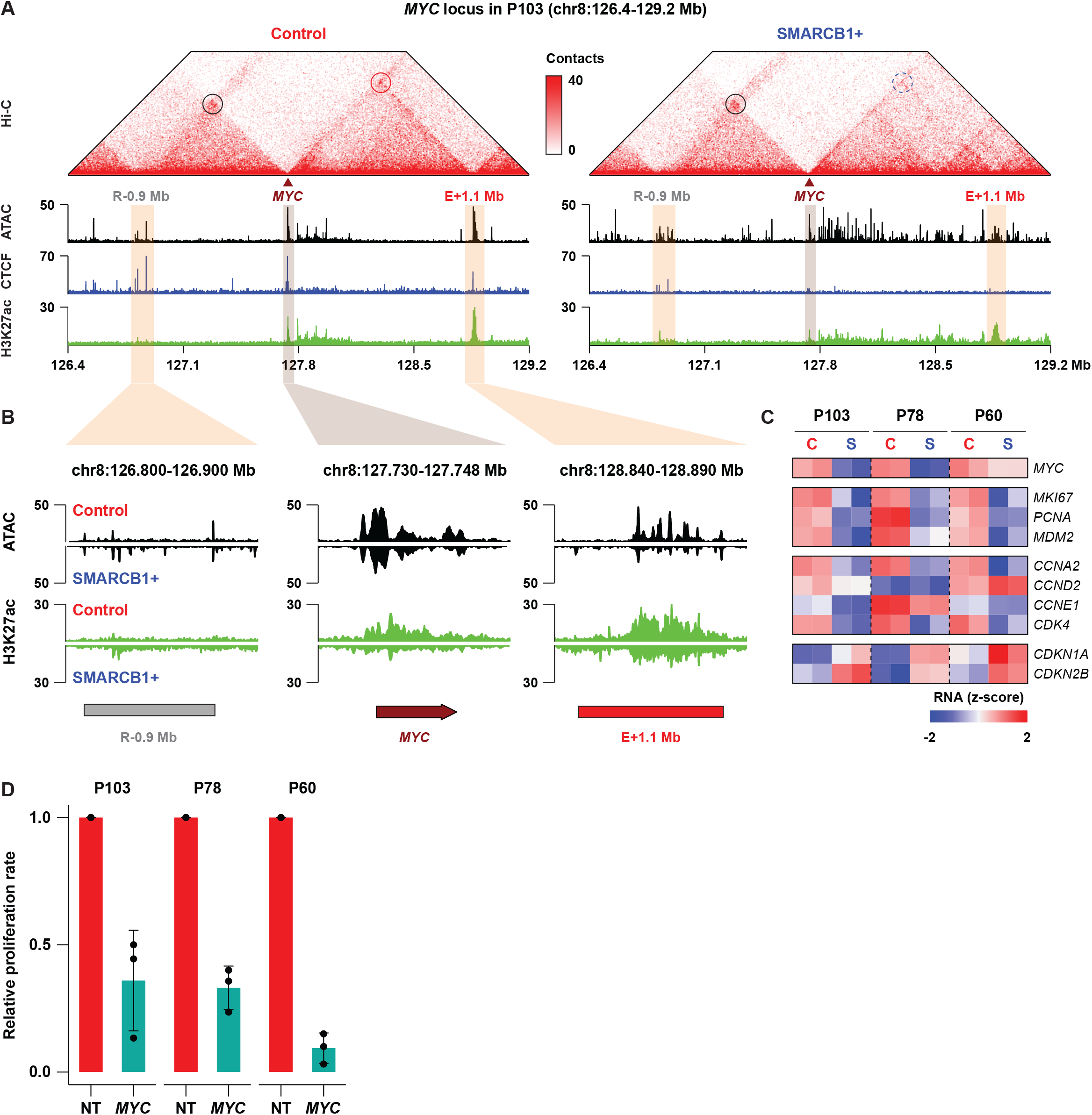
SMARCB1 controls a novel distal enhancer of the *MYC* oncogene in the MRT PDO model. (A) Hi-C analysis identifies two putative distal regulatory regions of *MYC* that interact with its promoter (n = 1); (B) A further epigenomic characterization suggests that the 1.1 Mb downstream regulatory element is a novel super enhancer of *MYC*, which can be robustly repressed by *SMARCB1* reconstitution; (C) *MYC* is consistently overexpressed in the *SMARCB1* mutated tumor PDOs derived from three different patients. Higher *MYC* level in these tumors is also associated with an increased expression of cell proliferation markers and deregulation of the *MYC*-target cell cycle genes; (D) Knockdown of *MYC* leads to a decreased proliferation of the tumor PDOs (n = 3).

**Fig. 3:**
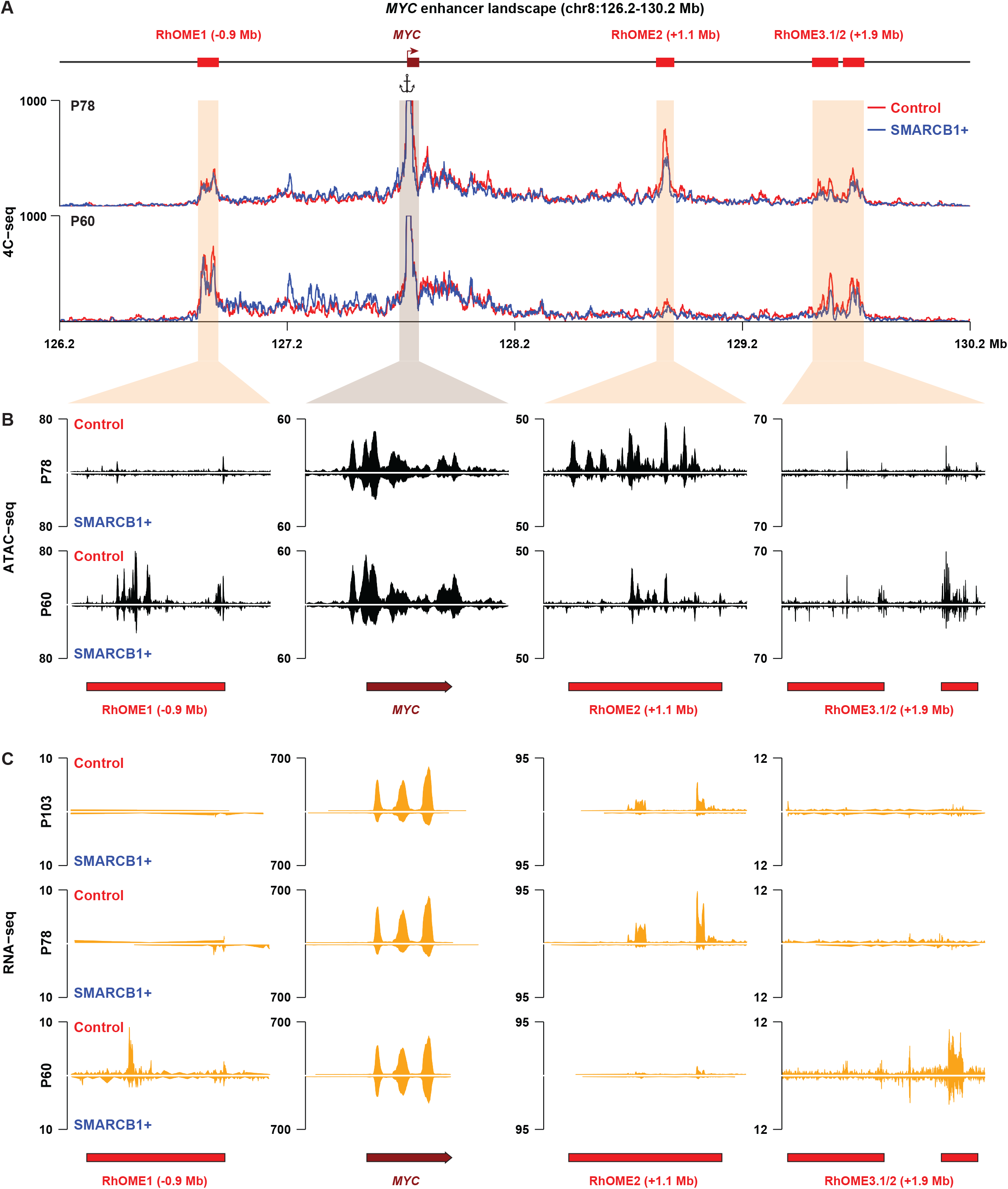
Identification of patients-specific *MYC* enhancer-ome in different MRT PDOs. (A) 4C-seq analysis identifies distinctive enhancer landscape in two other MRT PDO models (n = 2); (B) All the three distal *MYC* super enhancers can be repressed by *SMARCB1* reconstitution; (C) Expression of eRNAs indicates the activity of the *MYC* super enhancers.

The *MYC* oncogene is of particular interest in the context of MRT, as it was previously demonstrated by us and others that MRTs are defined by a SMARCB1-dependent *MYC* gene expression signature^8,23^. Similarly, we observed a strong loss of *MYC* expression as well as deregulation of cell cycle-related genes upon *SMARCB1* reconstitution, indicative of a cell cycle arrest (Fig. 2C). Moreover, shRNA-induced knockdown of *MYC* in MRT PDOs resulted in a significant reduction of proliferation (Fig. 2D and Extended Data Fig. 3E), indicating that MYC is required for MRT growth. In addition to a decrease in *MYC* expression, we observed that chromatin accessibility at the super enhancer as well as at the *MYC* promoter strongly diminished upon reconstitution of *SMARCB1* (Fig. 2A,B). These results show that SMARCB1 is required to prevent the formation of a distal super enhancer that interacts with the promoter of *MYC* through chromatin looping.

### Patient specific super enhancers interact with *MYC* in MRT

We hypothesized that *MYC* distal enhancer looping may be a general mechanism driving MRT oncogenesis. To this end, we performed *SMARCB1* reconstitution in two additional MRT PDOs (P78 and P60). RNA-seq analysis revealed a decrease in *MYC* expression in both these MRT models, as well as deregulation of cell cycle genes confirming the induction of a proliferation arrest (Fig. 2C,D). Next, we performed 4C-seq, a chromosome conformation capture method targeted to a specific genomic site, using the *MYC* promoter as the viewpoint, on these two additional PDOs. Similar to P103, P78 showed a reduction in *MYC* promoter contact frequency with the same ∼1.1 Mb distal genomic region (Fig. 3A) upon *SMARCB1* reconstitution. Remarkably, 4C-seq analysis in the third PDO model (P60) showed no interaction of the *MYC* promoter with our previously identified genomic region. Instead, 4C-seq uncovered two other chromatin loops that were diminished upon *SMARCB1* reconstitution (Fig. 3A).

To chart the regulatory landscape of the PDO models in the presence and absence of SMARCB1, we performed ATAC-seq. When we overlaid the ATAC-seq data with the 4C-seq data, we found that the regions that interact with the *MYC* promoter were highly accessible. The accessibility correlates with the patient-specific *MYC* promoter interaction landscape (Fig. 3B). Upon *SMARCB1* reconstitution, the loss of interactions coincides with a loss of accessibility at these loci (Fig. 3B). The stretches of accessible chromatin in P78 resemble the super enhancer we identified in P103. Collectively, we refer to these putative super enhancer regions as Rhabdoid Oncogenic *MYC* Enhancer (RhOME) and number them according to their position in the genome. Notably, in contrast to RhOME2, RhOME1 (*PCAT1*) and RhOME3 (*CCDC26*) were previously identified as *MYC* regulatory regions in other tumor entities^24–26^. Our Hi-C and 4C experiments thus revealed that in different MRT PDOs, distinct distal enhancers interacting with the *MYC* promoter are active (RhOME1-3). The intertumoral specificity of the RhOMEs was further highlighted by the expression of enhancer RNAs (eRNAs), distinctive for active super enhancers^27^, specifically in the MRT PDOs in which these enhancers are active and interacting with the *MYC* promoter (Fig. 3C). These results indicate that loss of *SMARCB1* induces an epigenetic state that is characterized by the formation of long-range promoter-enhancer loops that are specific to a PDO model, but heterogeneous between MRT PDOs derived from different donors.

Because MRT PDOs, to a large extent, retain the (epi-)genetic characteristics of the tissues they were derived from^17,28^, we used these models to further explore patient-specific epigenetic programs using ATAC-seq (Extended Data Fig. 4A,B). In addition to 29 and 556 OCRs that are consistently lost (Control-specific) or gained (SMARCB1+-specific) after *SMARCB1* reconstitution, respectively, we find many OCRs that are either specific for a single MRT PDO model or shared between two out of the three models (Extended Data Fig. 4B). In fact, 81.6%, 72.1% and 78.7% of the OCRs lost in P103, P78 and P60, respectively, are lost specifically in that PDO. In contrast, transcriptomic changes show less heterogeneity between the three MRT PDOs (Extended Data Fig. 4C). To stratify the OCRs that are lost in P103, we performed *k*-means clustering including the ATAC-seq data from the three PDOs and the ChIP-seq data from P103. Consequently, we could classify the lost OCRs into (K1) non-enhancer CTCF binding sites, (K2) weak enhancers, and (K3) CTCF-independent super enhancers (Fig. 4A and Extended Data Fig. 5A,B). Consistent with the function of super enhancers in control of cell identity^29^, we identified the SOX protein motifs, including SOX2, SOX9 and SOX17, and functional annotations of many developmental processes that are associated with the K3 cluster-specific open chromatin sites (Extended Data Fig. 5C,D). Whereas the K1 and K2 clusters show a similar accessibility pattern in all three PDO models, the accessilibity loss of the K3 cluster is specific to the P103 line (Extended Data Fig 5A). Our Hi-C analysis in P103 shows that these K3 cluster-specific super enhancers form insulation boundaries, which were essentially abolished upon reconstitution of *SMARCB1* (Fig. 4B, C). Thus, *SMARCB1* reconstitution in MRT organoids PDOs coincides with dramatic but local, patient-specific changes in genome topology and enhancer activity.

**Fig. 4:**
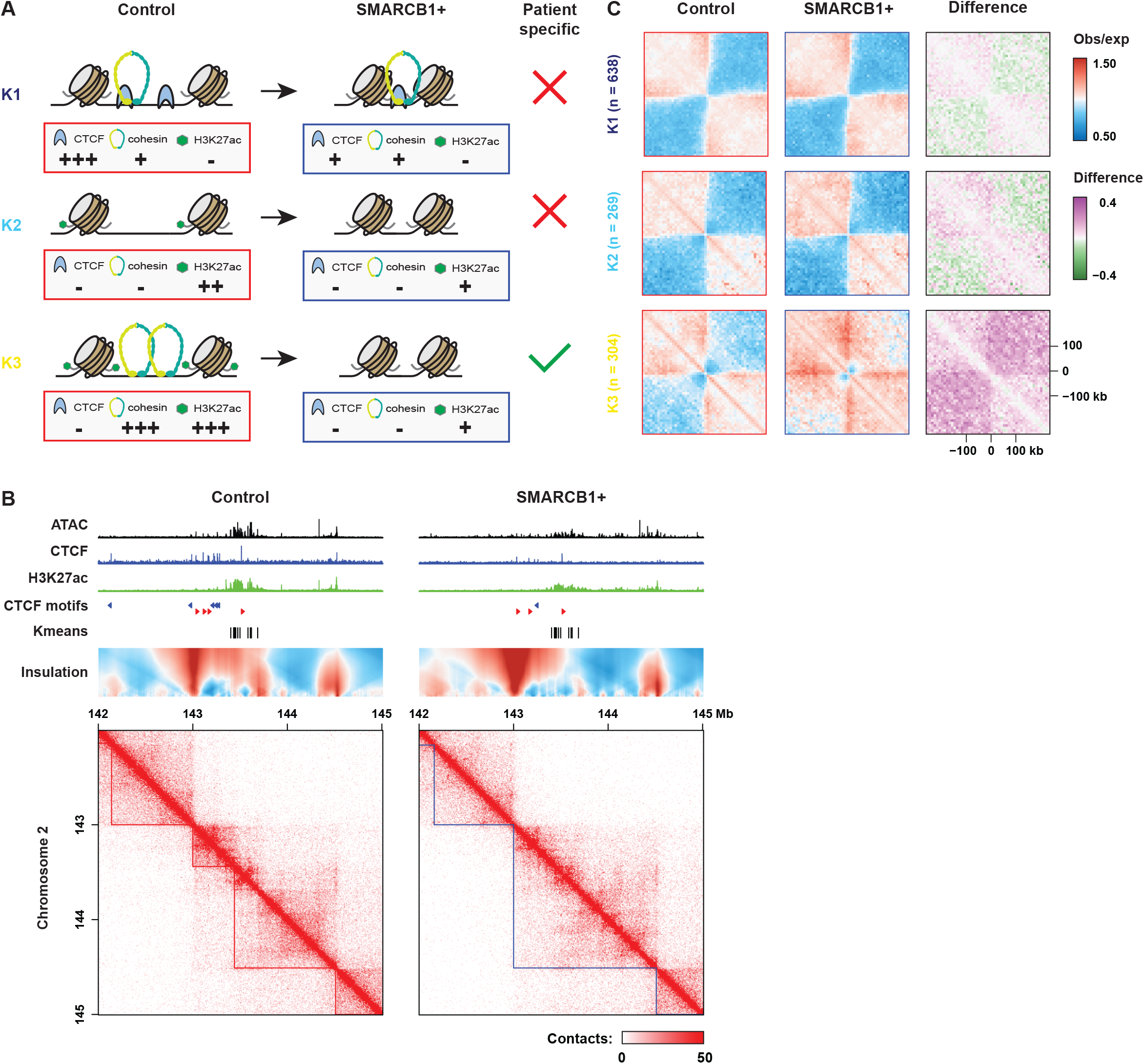
Patient-specific super enhancers form TAD boundaries in the SMARCB1-null tumor PDOs that are independent of CTCF. (A) A schematic model summarizes the three different types of control-specific open chromatin sites classified based their local epigenomic contexts; (B) A super enhancer forms a TAD boundary that is diminished after *SMARCB1* reconstitution; (C) Aggregate Region Analysis (ARA) shows that patient-specific super enhancers form SMARCB1-sensitive insulation borders in the tumor PDOs.

**Fig. 5:**
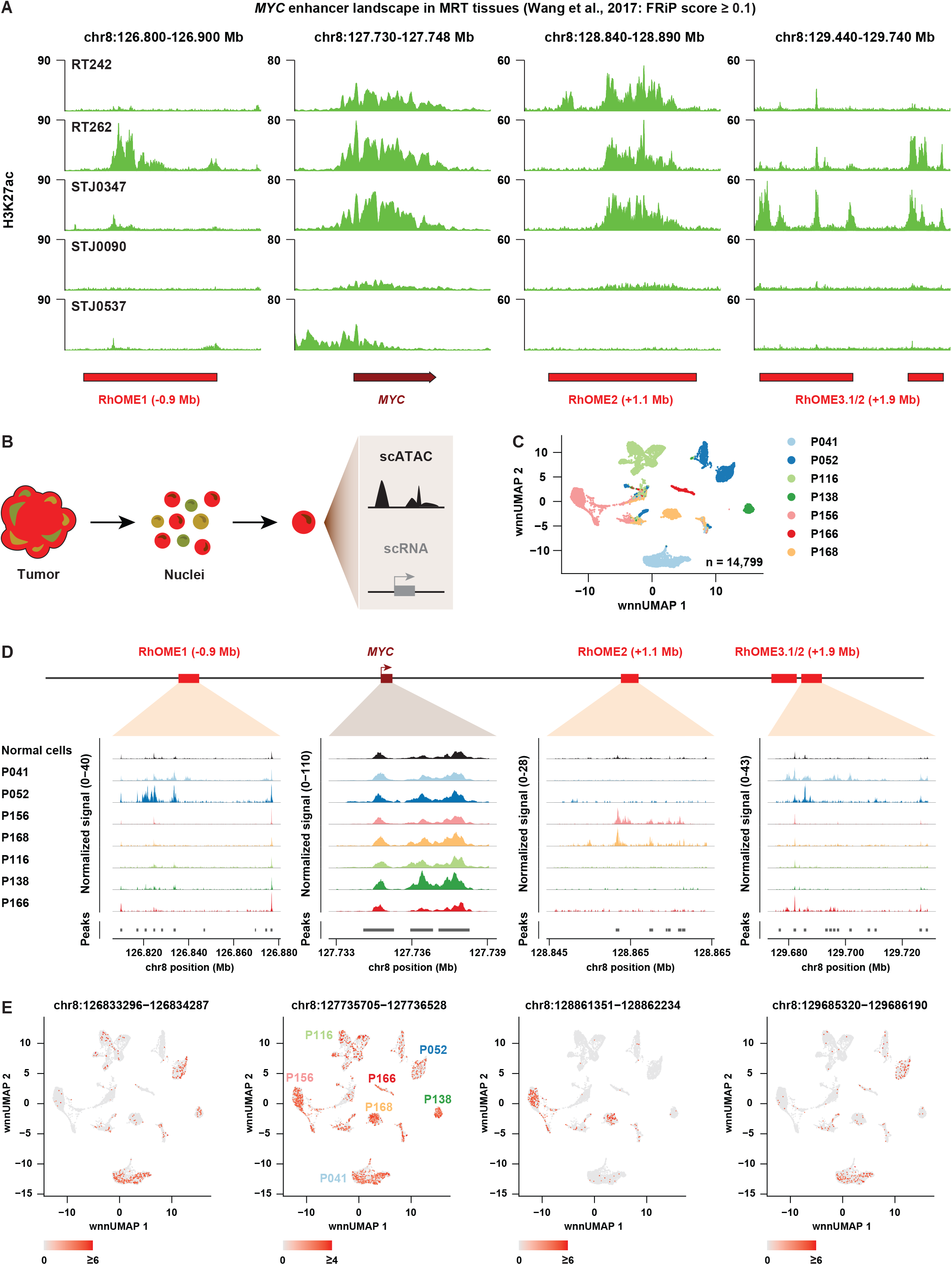
Analyses of MRT tissues reveal inter- and intra-tumor heterogeneity of *MYC* enhancer landscape. (A) The level of histone mark H3K27ac, an indicator of active promoters and enhancers, is coregulated at the *MYC* promoter and its super enhancers in MRT tumor tissues; (B) Acquisition of single-cell multiome data from MRT tissues using Chromium 10X Genomics platform; (C) UMAP discriminating tumor cells, but not normal cells, from the MRT tissues derived from different patients; (D) A pseudo-bulk view of scATAC data unravels inter-tumor heterogeneity of *MYC* regulatory landscape; (E) UMAP depicting intra-tumor heterogeneity of the *MYC* promoter and enhancers at single-cell level.

### *MYC* enhancer plasticity is reflected in patient MRT

Our data so far raise the intriguing possibility that there is intertumoral heterogeneity on the level of enhancer utilization driving expression of the *MYC* oncogene. To extrapolate our *in vitro* findings to patient tissues, we first examined publicly available H3K27ac ChIP-seq profiles from patient MRT tissues^30^. As expected, the *MYC* promoter is marked by high levels of H3K27ac in the majority of patient samples, indicative of active transcription (Fig. 5A). Moreover, broad enrichment for H3K27ac could be detected at RhOME1-3, showing that tumor super enhancer landscapes identified in MRT PDOs are preserved in MRT tissues. No peaks at RhOME1-3 could be detected in STJ0090 and STJ0537, which also show weak H3K27ac signals at the *MYC* promoter. Remarkably, and in line with our PDO data, the presence of H3K27ac peaks at the different RhOME loci was heterogeneously distributed across the other patient samples (Fig. 5A).

To assess whether differential enhancer utilization occurs within a tumor, i.e. intratumoral heterogeneity, we performed combined single-cell RNA-seq and ATAC-seq on seven MRT patient tissue samples using single-cell multiomics (Chromium 10X Genomics platform) (Fig. 5B). After filtering, 14,799 cells were left for further analyses, with a median of 2,040 cells per tumor (Fig. 5C). Normal and tumor cells were assigned by cell type marker gene^31^ and *SMARCB1* expression (Extended Data Fig. 6A-D). Uniform Manifold Approximation and Projection (UMAP) space revealed that tumor cells clustered by patient, while non-malignant cells clustered by cell type (Fig. 5C). *MYC* expression as well as OCRs at the *MYC* promoter were found across all MRTs and some normal cell clusters (Fig. 5D and Extended Data Fig. 6E). However, despite detectable *MYC* expression, almost no accessible chromatin could be detected at RhOME1-3 in normal cells (Fig. 5D), suggesting that these super enhancers are tumor specific in these samples. Crucially, we found different combinations of OCRs at RhOME1-3 in the different patient MRTs (Fig. 5D,E). More specifically, while P156 and P168 are exclusively defined by OCRs at RhOME2, P052 and P041 primarily show open chromatin regions at RhOME1 and RhOME3 and in P166 OCRs were only found at RhOME3. OCRs were not found at any of the RhOMEs in P116 and P138 suggesting supraphysiological activation of *MYC* occurs through regulatory elements other than RhOME1-3 in these tumors. Collectively, these results show for the first time the existence of intertumoral heterogeneity on the level of super enhancer activity driving expression of the *MYC* oncogene in MRT.

### ncBAF inhibition induces gene expression signatures that resemble *SMARCB1* reconstitution

It was recently demonstrated that MRT may be driven by the residual ncBAF complex aberrantly localizing at super enhancers in MRT cells^30,32,33^. As such, inhibition of the ncBAF subunit BRD9 was demonstrated to be a putative therapeutic vulnerability in MRT. We therefore set out to investigate the effect of ncBAF inhibition on MRT PDOs (Fig. 6A). To do so, we treated MRT PDOs with a pharmacological inhibitor of BRD9 (iBRD9)^32,33^. We observed that, morphologically, MRT cells exhibited a differentiation phenotype similar to *SMARCB1* reconstitution^8^ (Fig. 6A). Quantitative RT-PCR (RT-qPCR) confirmed that BRD9 inhibition caused a significant decrease in *MYC* mRNA levels (Fig. 6B). We performed RNA-seq to measure the transcriptomic changes following ncBAF inhibition. Visual inspection of the differentially expressed genes in *SMARCB1*+ cells suggests that the transcriptomic changes in the iBRD9-treated PDOs are similar (Fig. 6C). To statistically confirm the overlap between the differential gene sets, we performed a Fisher’s exact test on the two sets (Fig. 6D), which showed a strong similarity. Consistent with a decrease in proliferation, we observe a downregulation of cell cycle related gene sets as well as MYC target genes (Fig. 6E). In the upregulated gene sets we found an enrichment of genes associated with differentiation (Fig. 6E), consistent with the morphological phenotype of the cells treated with the BRD9 inhibitor (Fig. 6A). Thus, inhibition of the ncBAF complex in MRTs, eliminating all three BAF complexes, phenocopies the effects of *SMARCB1* reconstitution that restores all the BAF complexes.

**Fig. 6:**
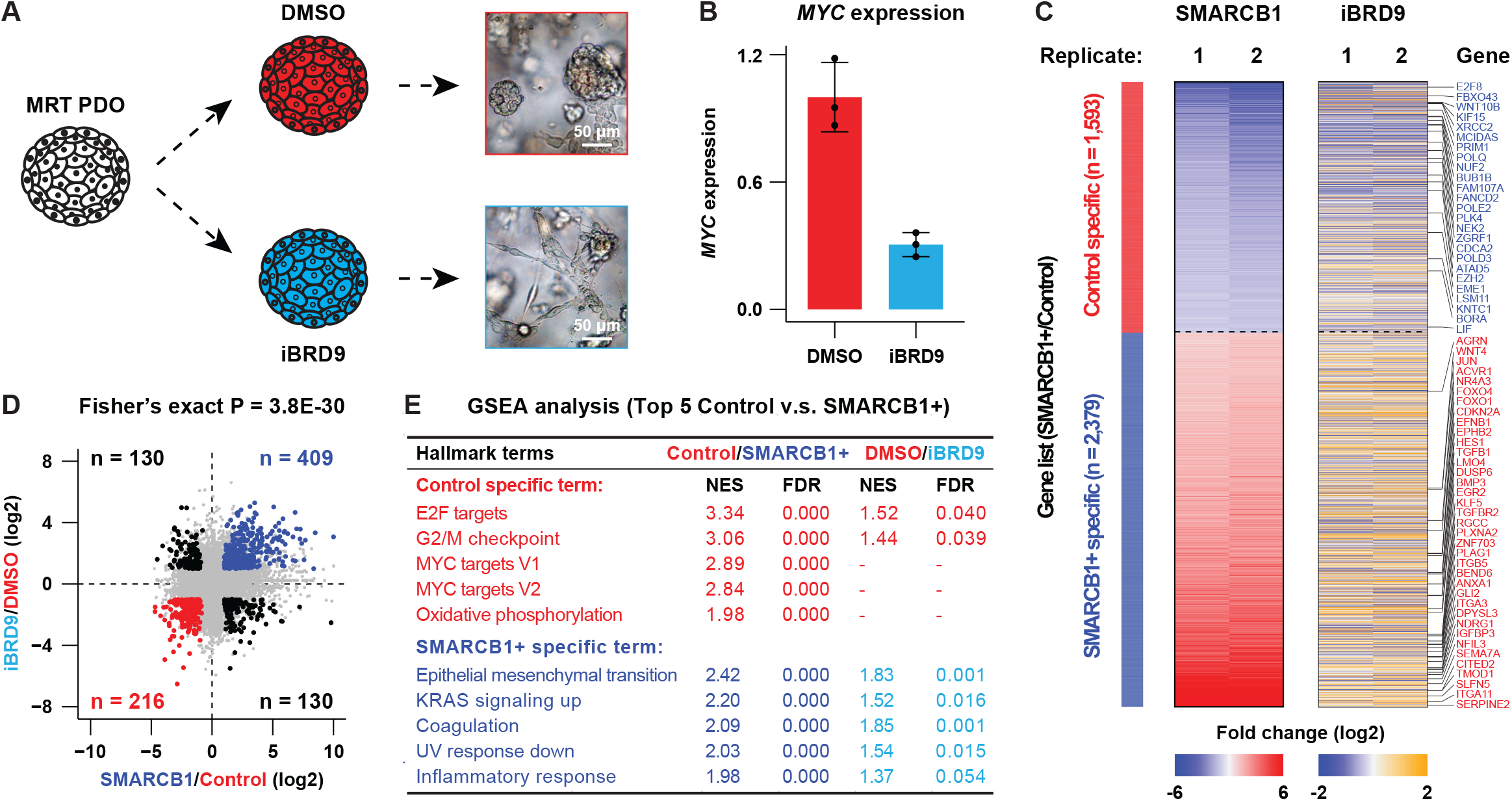
A therapeutic strategy for MRT patients via targeting a ncBAF subunit named BRD9 using a pharmacological inhibitor. (A) BRD9 inhibition leads to a differentiation phenotype of the MRT PDOs; (B) Expression of *MYC* is drastically reduced after BRD9 inhibition (n = 3); (C,D) *SMARCB1* reconstitution and BRD9 inhibition result in the similar transcriptomic changes (n = 2); (E) GSEA analyses reveal that many top hallmark gene sets identified in the *SMARCB1* reconstitution experiment are also differentially regulated after BRD9 inhibition.

**Fig. 7:**
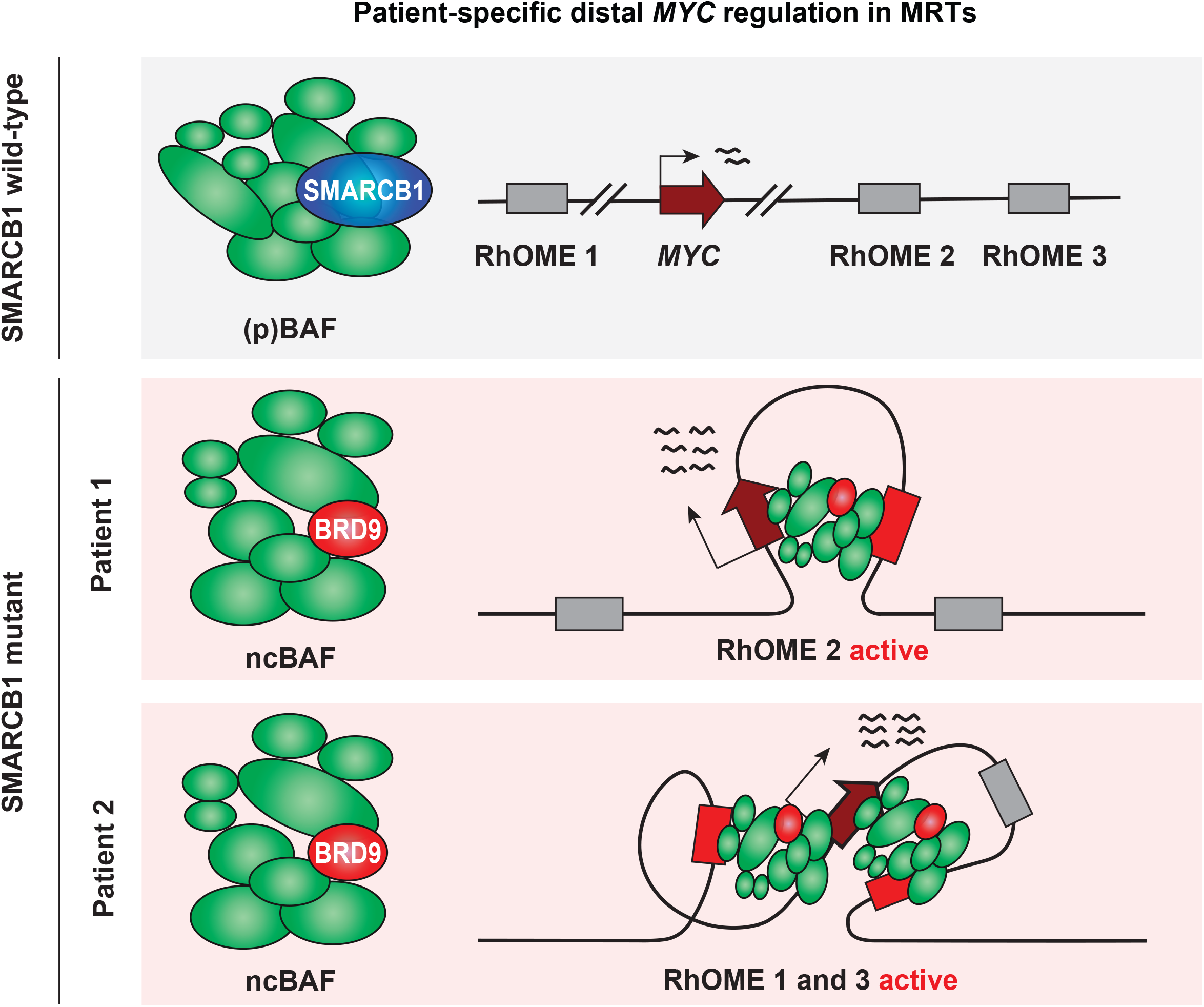
A graphic abstract summarizes *MYC* regulation in MRTs by distal patient-specific super enhancers.

Altogether, our comprehensive study provides the first evidence that derailed activity of super enhancers caused by *SMARCB1* loss underpins MRT tumorigenesis and serves as a blueprint for unraveling the contribution of BAF complex mutations to tumorigenesis across cancers.

## Discussion

Decades of cancer genetics research has revealed that adult tumors are unique based on their genetic profile, with each patient tumor having a distinctive mutation landscape^34,35^. Although adult cancers can harbor hundreds to thousands of mutations, recurrent driver mutations can be identified^36^. Typically, specific signaling pathways affecting cell proliferation and survival can be affected by mutations in different pathway members. Pediatric tumors on the other hand have a low mutational burden and are characterized by a small number or even a single driver event^37^. Additional epigenetic changes may contribute to pediatric tumor formation. Identifying recurrent patient-specific epigenetic driver landscapes has remained challenging. Because PDOs maintain many molecular characteristics of their parental tumors they can be used to study patient-specific changes on the epigenetic level as well. Our BAF complex reconstitution experiments enabled us to zoom into putative oncogenic enhancers. Upon analyzing the (single-cell) regulatory landscapes of primary MRT tissue we were able to identify potential super enhancers that were recurrently activated in a patient-specific manner.

A common denominator of MRT is the high expression of the *MYC* oncogene^8,23^. The regulatory mechanisms causing aberrant *MYC* activation in MRT has so far remained unknown. Here, we identify three putative super enhancers looping to the *MYC* promoter in MRT, as a consequence of *SMARCB1* loss. One of these super enhancers, RhOME2, has not been described before in the context of tumorigenesis. While different RhOMEs loop to the *MYC* promoter in different tumors (intertumoral heterogeneity), we find preferential use of one or a combination of RhOMEs within the same tumor. The patient-specific enhancer landscapes found in MRT could reflect the developmental origin of MRT, which lies in the neural crest^8^. Neural crest development is characterized by rapid switching of cell states caused by, amongst others, chromatin reorganization to assure quick and simultaneous development of several different cell types^38–40^. We hypothesize that loss of *SMARCB1* in a specific cellular context during neural crest development prevents the inactivation of certain *MYC* enhancers, which is essential for proper lineage specification. Ultimately, these *MYC* enhancers may transform to an oncogenic *MYC* super enhancer and drive rhabdoid tumorigenesis. Depending on the cellular identity of the tumor initiating cell, different enhancer landscapes may be active, possibly explaining why we find patient-specific enhancers. During development, cellular identity is in part determined by growth factors and morphogens that signal to transcription factors to establish a cell-type specific regulatory landscape. How this regulatory landscape is maintained in the absence of these signaling molecules is an interesting question for the future.

Our work reveals a role for SMARCB1 in 3D genome organization regulating long-range promoter-enhancer loops. Furthermore, SMARCB1 is required to prevent the formation of super enhancer-driven TAD boundaries, which are observed in MRT cells. In a previous study, restoration of SMARCB1 in an epithelioid sarcoma cell line showed hardly any effects on 3D genome organization. Although in these cells *MYC* interacts with the genomic region of RhOME3, no difference was observed upon SMARCB1 reconstitution^41^. We hypothesize that SMARCB1 executes its role in 3D genome organization by preventing the formation of ectopic super-enhancers which may be cell-type specific. Super enhancers are bound by cohesin and we also observed this in the MRT cells explaining how the very distal RhOMEs can interact with the *MYC* promoter. We and others have previously shown that accessible chromatin regions, which are directly regulated by the BAF complex, are important for cohesin positioning at active enhancers^21,42,43^. Therefore, *SMARCB1* inactivation may alter long-range chromatin interactions via reshaping cohesin occupancy in the mutant cells. *SMARCB1* reconstitution drastically weakened CTCF binding, but the CTCF-anchored chromatin loops and the majority of TAD boundaries remained mostly unaffected. Consistent with a model of cohesin-mediated loop extrusion we expect that cohesin is loaded at the distal super enhancers and brings the enhancer and the promoter in close proximity. The reported enhancer-docking CTCF site in the *MYC* promoter^44^ likely plays a role in the interactions between the *MYC* promoter and the RhOMEs. To further understand enhancer selection in MRTs, a mechanistic study is required to delineate the regulome for each of the *MYC*-RhOME loops.

By inhibiting BRD9, and therefore repressing the activity of the non-canonical BAF complex, we could show that loss of the ncBAF complex has a similar effect as rescuing the cBAF complex. Therefore, MRT initiation during development might be dependent on the activity of the ncBAF complex. In addition to MRTs, several other pediatric cancers are characterized by aberrations in the BAF complex^5^. Moreover, BAF complex members are among the most commonly mutated genes in adult malignancies^4^. Using our study as a blueprint for other BAF complex-mutant cancers could shed light on the importance of intertumoral epigenetic heterogeneity to tumorigenesis and the therapeutic potential of ncBAF targeting on a broader scale.

## Acknowledgements

We acknowledge funding from the European Research Council (ERC) (ERC-StG #850571 to J.D.), the Foundation Children Cancer Free (KiKa 338, L.C. and I.P.), and the Dutch Research Council (NWO-Vidi 203.003 (J.D.). Work in the de Wit laboratory is supported by the European Research Council (ERC) (ERC-CoG #865459) and the Dutch Research Council (NWO-Vidi, ‘016.16.316’). N.Q.L. is supported by a Veni grant from the Netherlands Scientific Organization (NWO, 016.Veni.181.014). Additional support was provided by the American Association for Cancer Research (AACR) and the St. Baldrick’s Foundation (Pediatric Cancer Research Award to J.D.). We thank the NKI Genomics Core Facility and Princess Máxima Center Single Cell Genomics Facility for technical support, as well as the Princess Máxima Center Biobank and Data Access Committee for providing MRT tissues. We are profoundly grateful to the patients and parents who agreed to participate in our research.

## Methods

### Ethics statement

MRT tissues used for single-cell multiomics (Supplementary Table 1) were obtained under approval by the medical ethical committee of the Erasmus Medical Center (Rotterdam, the Netherlands) and the Princess Máxima Center for pediatric oncology (Utrecht, the Netherlands). Written informed consent was provided by all patients and/or parents/guardians.

### Patient-derived organoid culture

MRT PDOs were cultured as previously described^17^. For BRD9 inhibition experiments, MRT PDOs were mechanically dissociated and plated at a density of 10.000 cells/ul in 10ul BME droplets. After 24h, 10μM I-BRD9 inhibitor (MedChemExpress, HY-18975) was added to the PDOs. PDOs were harvested 120h later for RNA isolation.

### Vector construction

pLKO.1-UbC-luciferase-blast and pLKO.1-UbC-hSMARCB1-blast lentiviruses were previously described^17^. For *MYC* knockdown experiments, the following targeting sequence was used: 5’-CCCAAGGTAGTTATCCTTAAA-3’.

### Lentiviral transductions

Lentiviral transductions were performed as described^45^. For *SMARCB1* re-expression, MRT PDOs were transduced with pLKO.1-UbC-luciferase-blast or pLKO.1-UbC-hSMARCB1-blast lentiviruses, as described^8^. After two days, 10 μg/ml blasticidin was added to the culture medium. MRT cells were harvested four days after transduction for the different applications. For *MYC* knockdown experiments, MRT PDOs were transduced with pLKO.1-puro lentiviruses. For each condition, 30,000 cells were seeded. After two days, puromycin was added to the culture medium. Nine days after lentiviral infection, PDOs were harvested for RNA extraction.

### Quantitative RT-PCR

PDOs were harvested in RA1 buffer and RNA was subsequently isolated using the Macherey-Nagel RNA isolation kit following the manufacturer’s instructions. The extracted RNA was used for cDNA production using GoScript reverse transcriptase (Promega). Quantitative RT-PCR was performed using IQ SYBR green mix (Biorad) following manufacturer’s instructions. Results were calculated using the ΔΔCt method. Primer sequences: *MYC*_Fw: 5’-GATTCTCTGCTCTCCTCGACG-3’, *MYC*_Rv: 5’-GATGTGTGGAGACGTGGCA-3’, *GAPDH*_Fw 5’-TGCACCACCAACTGCTTAGC-3’, *GAPDH*_Rv 5’-GGCATGGACTGTGGTCATGAG-3’.

### RNA-sequencing

RNA was isolated from the harvested PDOs using TRIzol RNA isolation reagent (Invitrogen) according to the manufacturer’s instructions. RNA-seq libraries were prepared using the TruSeq Stranded RNA LT Kit (Illumina). The libraries were sequenced on an Illumina NextSeq 550 platform using the paired-end 44+32 (75 cycles) mode. Two biological replicates were generated for each of the RNA-seq experiments in this study.

RNA-seq data were mapped against the hg38 reference genome by TopHat2 (version 2.1.1)^46^. Mapped reads with a mapping quality score <10 were removed using SAMtools. The read coverage of exons for each gene in the Homo Sapiens GRCm38.92 annotation file was determined with the HTSeq tool (version 0.9.1)^47^. The coverage files were generated with the ‘normalize to 1× genome coverage’ methods in deepTools. For visualization, we combined the two biological replicates of each condition to generate one merged bigWig file. The filtered coverage data was normalized using DESeq2 (version 1.24.0) with default parameters^48^. Differential peaks were detected using a Wald test (FDR <0.05 and fold change ≥2). Gene set enrichment analysis was performed with a desktop version of the GSEA tool (version 4.1.0)^49^ and the Molecular Signatures Database (MSigDB; V2022.1)^50^.

### ATAC-sequencing

PDOs were washed in ice-cold AdDF+++ and viably frozen in Recovery Cell Culture Freezing Medium (Thermo Fisher). For library preparation, cells were thawed and processed following an established protocol. In short, nuclei were isolated from cells and permeabilized. The isolated nuclei were tagmented using Tn5 transposase (Illumina, Nextera DNA Library Preparation Kit), followed by two sequential 9-cycle PCR amplification steps. The resulting DNA fragments (< 700 bp) were purified using SPRI beads (Beckman). ATAC-seq libraries were sequenced on a HiSeq 2500 (Illumina). For each of the ATAC-seq experiments, the data were recorded in biological triplicates.

ATAC-seq data were analyzed as previously described. In short, sequencing reads were mapped to the hg38 reference genome using BWA-MEM (version 0.7.15-r1140)^51^. The mapped reads were filtered using SAMtools^52^, discarding reads with mapping quality score <15, as well as optical PCR duplicates. The coverage files were produced using the deepTools (version 3.0) method “normalize to 1X genome coverage”^53^. For visualization, we combined the biological triplicates of each condition to generate one merged bigWig file. A merged peak list was generated from ATAC-seq data of MRT control and *SMARCB1*+ cells (n=3 independent experiments). The read coverage under the peaks was determined using a HTSeq tool (version 0.9.1)^47^. The peaks with ≥10 reads in each replicate were included for further analysis. The filtered coverage data was normalized using DESeq2 (version 1.24.0) with default parameters^48^. Differential peaks were detected using a Wald test (FDR <0.05 and fold change X2). Functional annotation of the differential peaks was performed using the GREAT analysis tool (version 4.0.4)^54^.

### ChIP-sequencing

The ChIP-seq experiments were performed according to an established protocol^21^. PDOs were washed in ice-cold AdDF+++ and PBS. Subsequently, cells were cross-linked with a final concentration of 1% formaldehyde for 10 min. Glycine (2.0 M) was used to quench the cross-linking reaction. The cross-linked cells were then lysed and sonicated using Bioruptor Plus sonication device (Diagenode) to obtain ∼300 bp chromatin. For ChIP, sonicated chromatin was incubated overnight at 4°C with antibodies that had first been coupled to Protein G beads (Thermo Fisher). After incubation, captured chromatin was washed, eluted and de-crosslinked. The released DNA fragments were purified using MiniElute PCR Purification Kit (Qiagen). The ChIP experiments were performed using the following antibodies: CTCF (07-729, Merck Millipore, 5 ml per ChIP), RAD21 (ab154769, Abcam, 2.2 mg per ChIP), and H3K27ac (ab4729, Abcam, 5 mg per ChIP). The purified DNA fragments were prepared using the KAPA HTP Library Preparation Kit (Roche) following manufacturer’s instructions. The libraries were sequenced on an Illumina HiSeq 2500 using the single-end 65-cycle mode.

ChIP-seq data were analyzed as previously described. In short, sequencing reads were mapped to the hg38 reference genome using the Bowtie 2 mapper (version 2.3.4.1)^55^. The mapped reads were filtered using SAMtools, discarding reads with mapping quality score <15, as well as optical PCR duplicates. The coverage files were produced using the deepTools (version 3.0) method “normalize to 1X genome coverage”^53^.

### Hi-C/4C

We generated Hi-C data as previously described with minor modifications^21^. In short, MRT PDOs (∼10 millions cells) were washed in ice-cold AdDF+++ and PBS. Subsequently, cells were cross-linked with a final concentration of 2% formaldehyde for 10 min. Glycine (2.0 M) was used to quench the cross-linking reaction. The restriction enzyme MboI was used to digest crosslinked DNA in the nucleus. At the restriction overhangs, biotinylated nucleotides were incorporated. Subsequently, overhangs were joined by blunt-end ligation. The ligated DNA was enriched by streptavidin pull-down. A standard end-repair and A-tailing method was used to further prepare the Hi-C libraries, which were sequenced on an Illumina Nova-Seq platform generating paired-end 150 bp reads.

Hi-C sequencing reads were processed using HiC-Pro^56^, which includes mapping, identification of valid Hi-C pairs, generation of contact matrices and ICE normalization. HiCCUPS (version 0.9) was used to call chromatin loops. Subsequent analyses were performed in GENOVA, a visualization tool of Hi-C data written in R^57^.

4C was performed as previously described^21^. In short, we used MboI as the first and Csp6I as the second restriction enzyme. The viewpoint was designed at the *MYC* promoter region using the primer pair: “CTCTTTCCCTACACGACGCTCTTCCGATCTTCTCCCTGGGACTCTTGATC” and “ACTGGAGTTCAGACGTGTGCTCTTCCGATCTGTCTGTTTAGCCCTGAGATG”. The 4C-seq libraries were sequenced using a NextSeq 550. Two biological replicates were measured for each of the 4C experiments in this study.

Mapping of the sequencing reads was performed according to our 4C mapping pipeline (http://github.com/deWitLab/4C_mapping). 4C data was normalized to 1 million intrachromosomal reads using peakC^58^. For visualization, we combined the two biological replicates.

### Motif analysis

Motif enrichment was computed using a similar method described in our previous publication^21^. We first identified and quantified the number of motifs for the peaks specific to MRT control or *SMARCB1*+ samples using GimmeMotifs (version 0.13.1) and the non-redundant cis-bp database (version 3.0)^59^. As a background peak set, we used the peaks that were unchanged upon *SMARCB1* re-expression. First, we normalized motif frequencies to the total number of identified motifs in that sample. Then, we calculated the log2-enrichment score of MRT control or *SMARCB1+* motifs by comparison to motif frequency in the background peak set. The p-value was calculated using the Fisher’s exact test.

### Single-cell multiomics

MRT tissues were processed using standard tissue processing procedure following the 10X Genomics protocol. In brief, tissues were minced into small pieces and homogenized using a dounce tissue grinder. After cell lysis, the sample was filtered once using a 70uM filter as well as a 40uM filter. Intact single nuclei were sorted and two independent samples were mixed based on different gender (male with female). Between 2.000 to 40.000 nuclei were loaded on the Chip J Chromatin Controller (10X Genomics). Library preparation of Gene expression (GEX) as well as ATAC library was performed following manufacturer instructions (10X Genomics). Initial processing of raw data files was done by cell ranger-arc (version 2.0.0), seurat (version 4.1.1)^60^ and signac (version 1.7.0)^61^.

Reads were mapped to GRCh38 and cell genotypes were annotated by souporcell^62^. Cell ranger-arc aggr function was used to harmonize detection of peak calling. Further filtering steps were included to filter cells of good quality (ATAC counts < 50000 & > 800, RNA counts < 30000 & > 800, nucleosome signal < 1.5, TSS enrichment > 1, % of blacklist regions < 3%, % of mitochondrial genes < 20%). Data was normalized using SCT normalization for RNA-seq data and TF-IDF normalization for ATAC-seq data. Gene marker expression (SingleR version 1.10)^63^ was used to annotate normal cell genotypes compared to tumor cells. Human Primary Cell Atlas Data was used as cell type reference (celldex version 1.6.0)^31,63^ for normal cell annotation. Normal versus tumor cell identification was verified by the absence of *SMARCB1* gene expression. UMAP plots were generated via joined clustering of both datasets (PCA 1:25, LSI 2:30).

## Data availability

RNA-seq, ATAC-seq, ChIP-seq and HiC data can be accessed via GSE218115 and combined single-cell RNA and ATAC sequencing data were deposited under GSE218385. The publicly available H3K27ac ChIP-seq data were downloaded using the accession number GSE71506, and reanalyzed using in house ChIP-seq pipeline as described in the “Methods” section.

## Figure legends

**Extended Data Fig. 1: A detailed characterization of differential open chromatin sites the MRT PDOs**. (A) *SMARCB1* reconstitution reprograms thousands of open chromatin sites in the MRT PDOs (n = 3); (B) Identification of transcription factor motifs that are enriched in the differentially accessible regulatory elements; (C) Top 5 most significant GO terms associated with the control and *SMARCB1+* MRT PDOs.

**Extended Data Fig. 2: Impact of *SMARCB1* reconstitution on higher-order of genome organization**. (A) *SMARCB1* reconstitution displays a mild effect on relative contact probability in the Hi-C analyses (n = 3); (B) An example locus shows that TADs are largely unaffected after *SMARCB1* reconstitution; (C) A total number of TADs identified in the control and *SMARCB1*+ PDOs; (D) Aggregate TAD analysis suggests that *SMARCB1* reconstitution only has weak effects on chromatin contacts within TADs; (E) Focal changes of A/B compartments after *SMARCB1* reconstitution; (F,G) In contrast to the control cells, the *SMARCB1*+ cells show a higher frequency of B compartment interactions and a lower frequency of A compartment interactions.

**Extended Data Fig. 3: SMARCB1 dependent looping preference in MRT cells**. (A) A total number of chromatin loops identified in the control and *SMARCB1*+ cells; (B) A waterfall plot summarizes changes of the identified chromatin loops before and after *SMARCB1* reconstitution; (C) The control-specific loops are significantly longer than other loops; (D) Different types of the identified loops based on their sensitivity to *SMARCB1* reconstitution; (E) Relative *MYC* expression measured by RT-qPCR in the three organoid lines following shRNA knock-down (n = 1).

**Extended Data Fig. 4: The response of open chromatin landscape and transcriptome to *SMARCB1* reconstitution in the three organoid lines**. (A) PCA analyses show that *SMARCB1* reconstitution leads to both consistent and patient-specific changes in chromatin accessibility and gene expression (ATAC-seq: n = 3, RNA-seq: n = 2. The PCA plots show an average of all the replicates); (B) Venn diagram analyses show the common and patient-specific changes of open chromatin landscapes after *SMARCB1* reconstitution; (C) The same analyses as in (B) but for transcriptomic changes.

**Extended Data Fig. 5: Molecular stratification of the common and patient-specific super enhancers in the MRT PDOs**. (A) A Kmeans clustering analysis identifies three groups of the control-specific (lost) open chromatin sites based on their molecular features in these MRT organoid lines; (B) Ranking H3K27ac signals of the three Kmeans clusters suggests that the majority of the K3 open chromatin sites are super enhancers; (C) Identifying transcription factor motifs that are overrepresented in the three types of open chromatin sites; (D) The top GO terms identified in each of the Kmeans clusters.

**Extended Data Fig. 6: Identification of patient-specific molecular groups using single-cell multi-omics**. (A) Single-cell multi-omics analysis identifies fifteen distinctive molecular groups from seven MRT tumor tissues; (B,C) UMAP space and hierarchical clustering analysis discriminate different cell types within these tumor tissues; The expression levels of (D) *SMARCB1* and (E) *MYC* in the fifteen molecular groups are visualized at single-cell level.

## Supplementary table

**Supplementary Table 1:** MRT tissues used for single-cell multiomics analysis.

| Sample | M-number | Male | Location | Category |
| --- | --- | --- | --- | --- |
| P052 | MRT_M711AAA_T | Male | Liver | Primary |
| P116 | MRT_M981AAC_T | Male | Liver | Primary |
| P166 | MRT_M728AAD_T | Male | Liver | Primary |
| P168 | MRT_M378AAD_T | Female | Vertebra | Primary |
| P156 | MRT_M375AAD_T | Female | Arm | Primary |
| P041 | MRT_M563AAB_T | Male | Ocular | Primary |
| P138 | MRT_M221AAD_T | Female | Liver | Primary |

