## Extended Data Figure 1 for "SMARCB1 loss creates patient-specific *MYC* topologies that drive malignant rhabdoid tumor growth"

### Extended Data Fig. 1

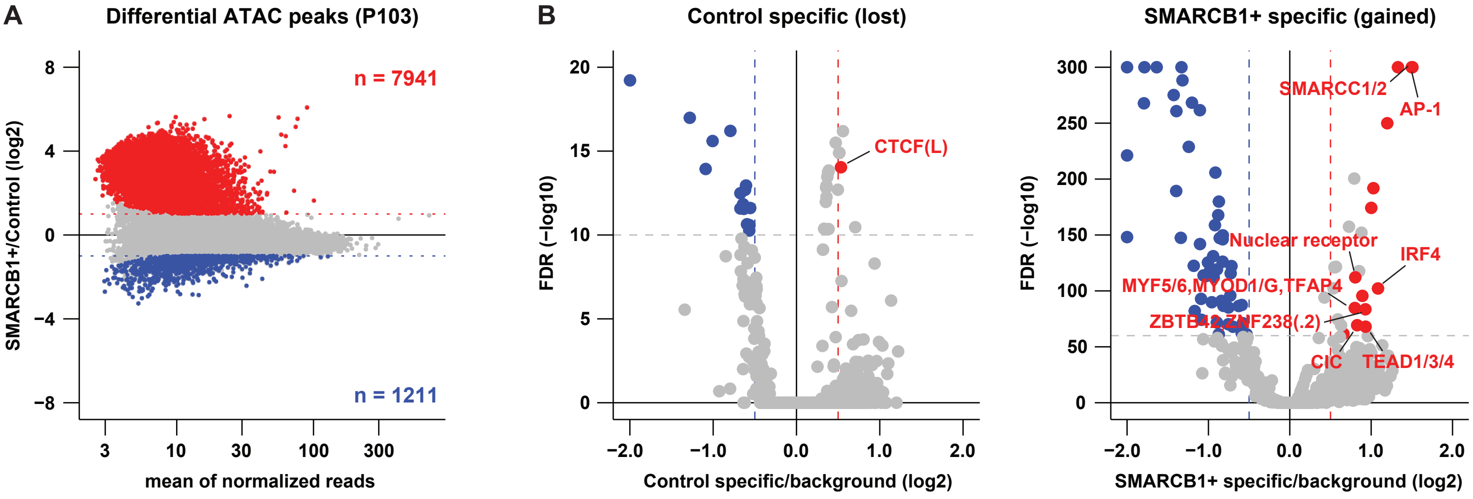


**C** Top 5 most significant biological processes (P103)

| ID | Biological process | Fold change | FDR |
|---|---|---|---|
| **Control specific (lost) term:** | | | |
| GO:0050777 | negative regulation of immune response | 2.61 | 1.08E-04 |
| GO:0045814 | negative regulation of gene expression, epigenetic | 2.71 | 1.89E-04 |
| GO:0006334 | nucleosome assembly | 2.43 | 8.16E-04 |
| GO:0050907 | detection of chemical stimulus involved in sensory perception | 3.27 | 9.17E-04 |
| GO:0043112 | receptor metabolic process | 2.52 | 9.99E-04 |
| **SMARCB1+ specific (gained) term:** | | | |
| GO:0072273 | metanephric nephron morphogenesis | 2.16 | 1.25E-13 |
| GO:0003338 | metanephros morphogenesis | 2.03 | 1.90E-13 |
| GO:0048643 | positive regulation of skeletal muscle tissue development | 2.17 | 3.14E-10 |
| GO:0072207 | metanephric epithelium development | 2.09 | 3.17E-10 |
| GO:0072170 | metanephric tubule development | 2.13 | 5.77E-10 |
