## Supplementary figures and images for "SMARCB1 loss creates patient-specific *MYC* topologies that drive malignant rhabdoid tumor growth"

### Extended Data Figure 2

#### Extended Data Fig. 2

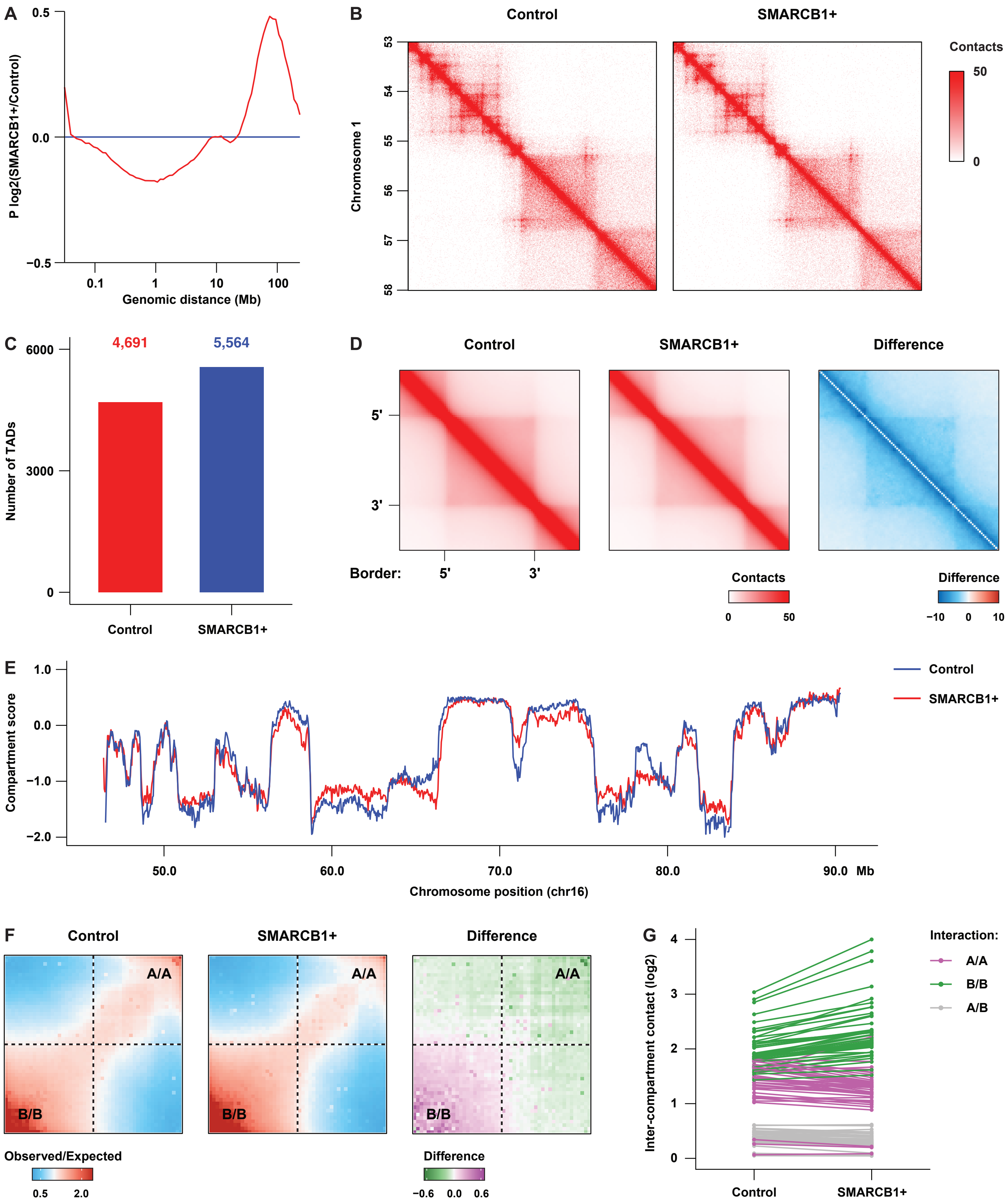

### Extended Data Figure 3

#### Extended Data Fig. 3

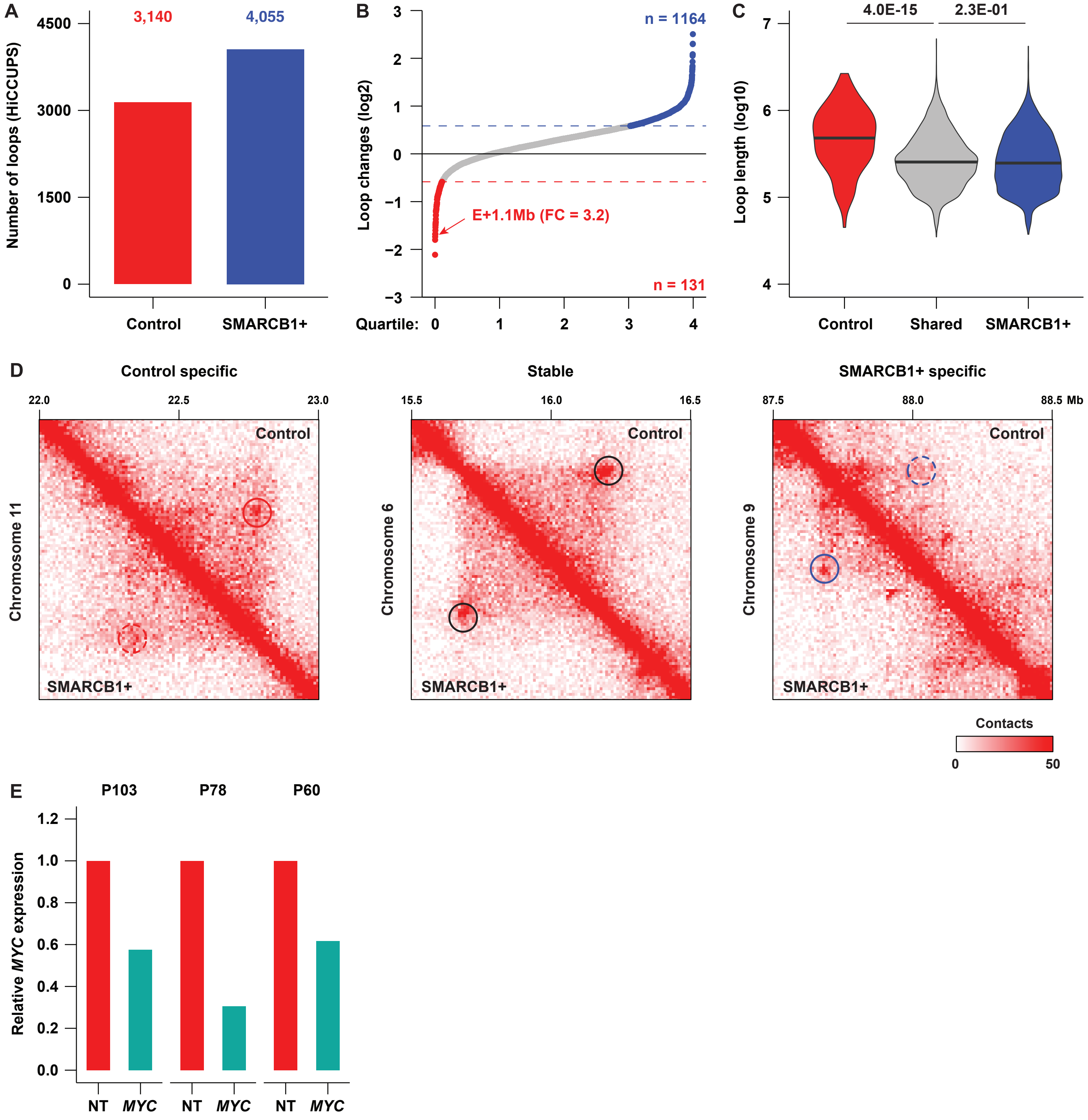

### Extended Data Figure 4

#### Extended Data Fig. 4

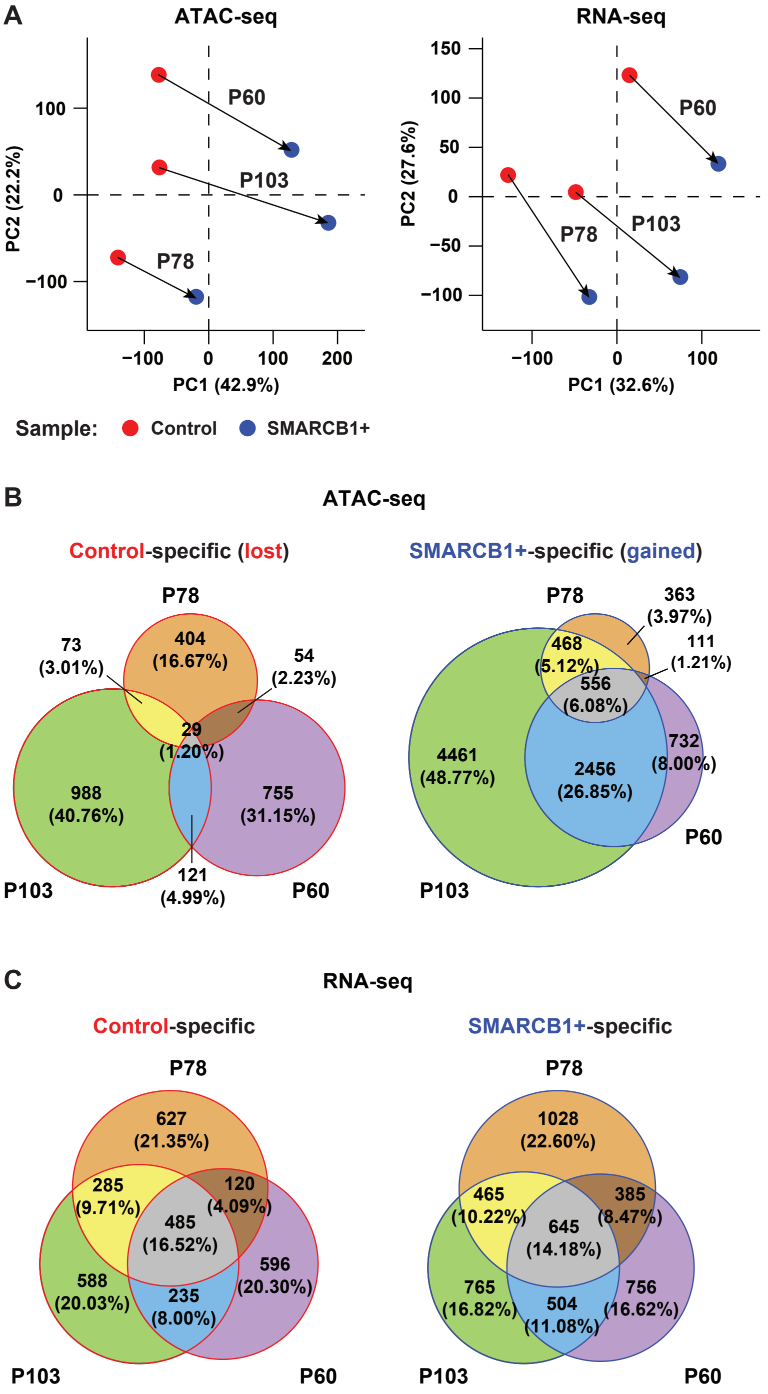

### Extended Data Figure 6

#### Extended Data Fig. 6

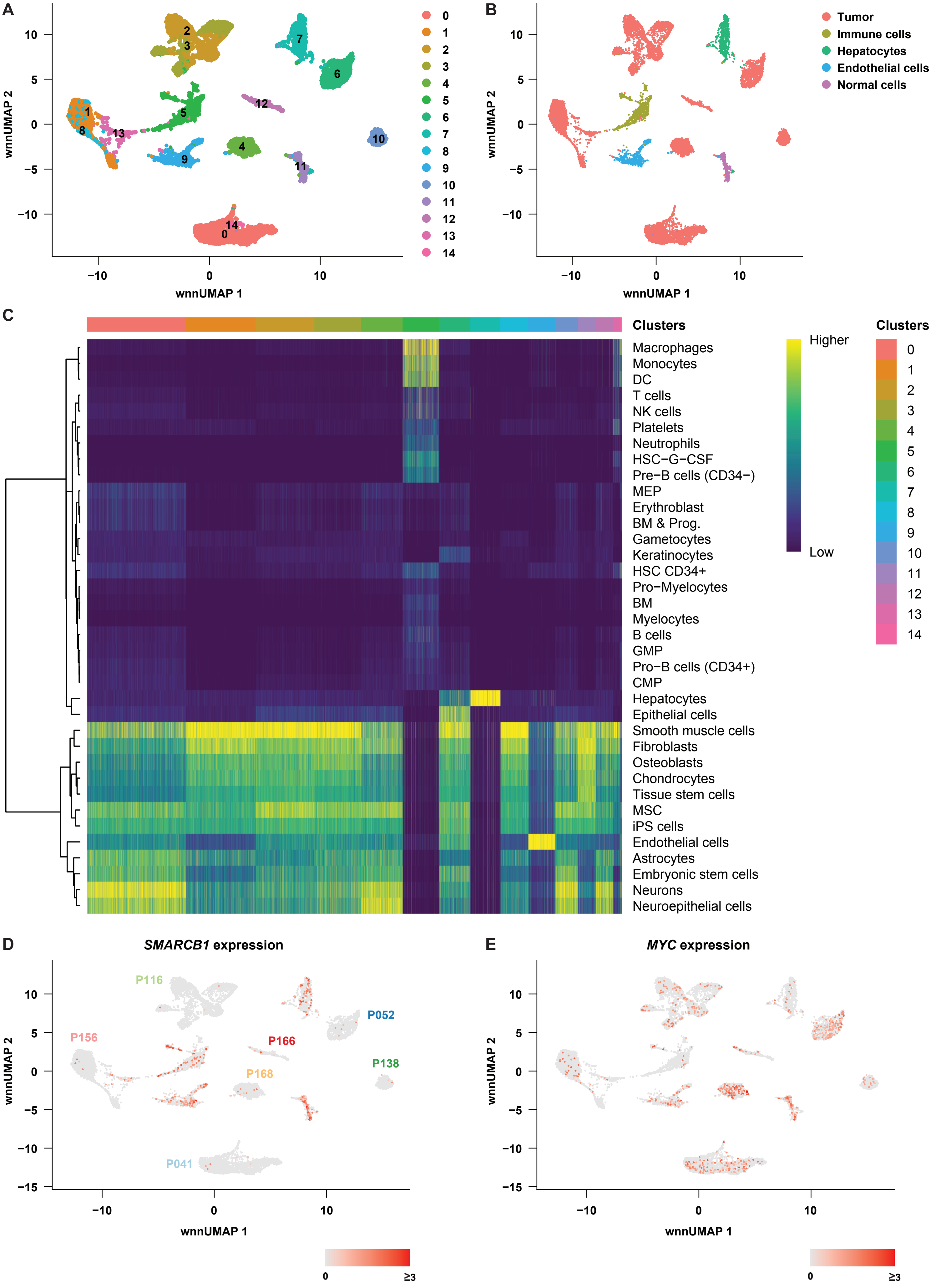
