## Extended Data Figure 5 for "SMARCB1 loss creates patient-specific *MYC* topologies that drive malignant rhabdoid tumor growth"

### Extended Data Fig. 5

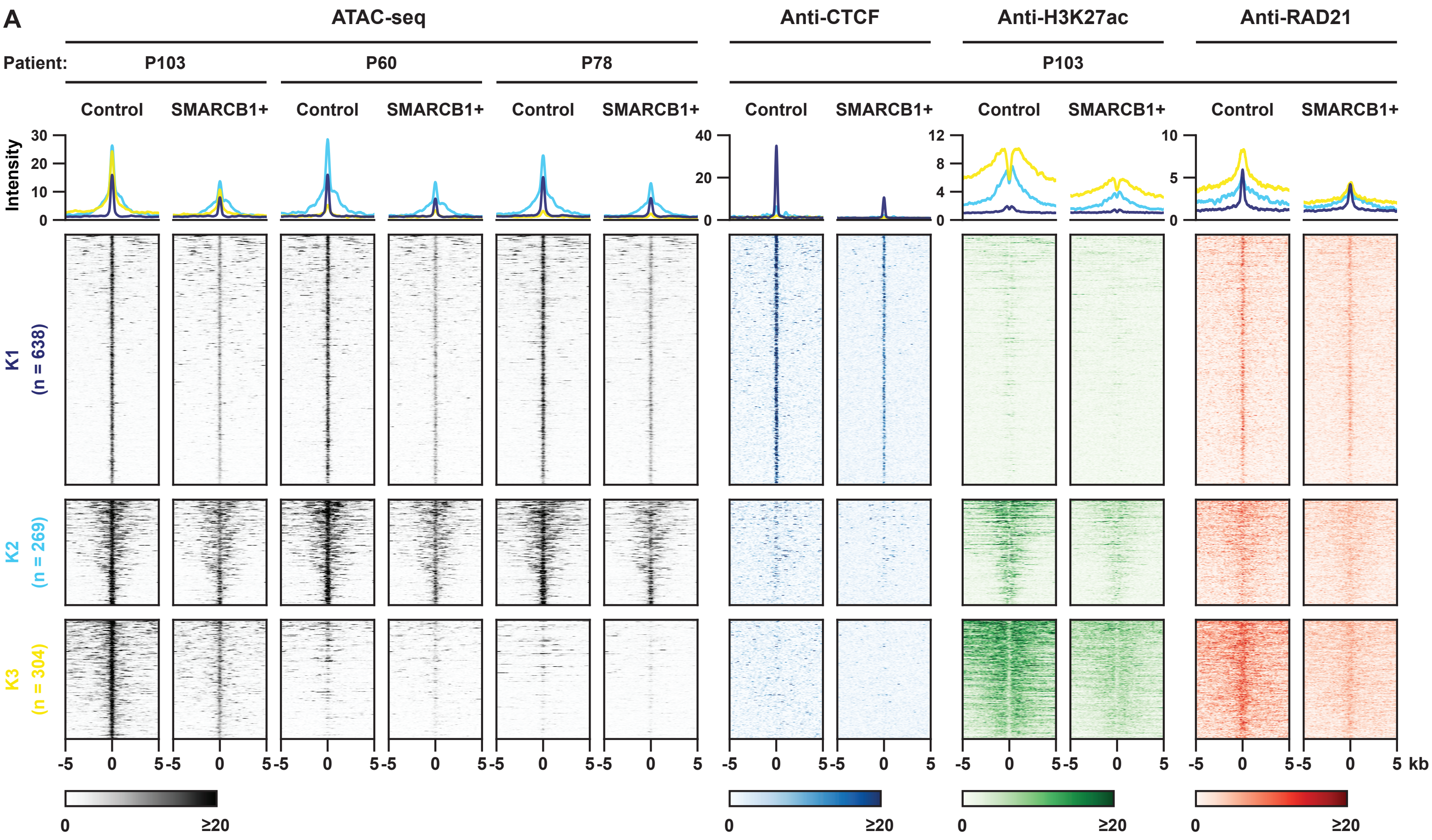


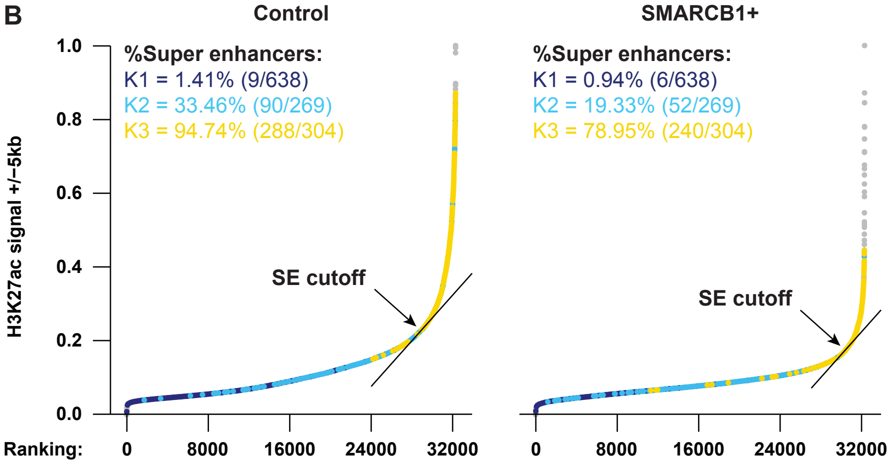


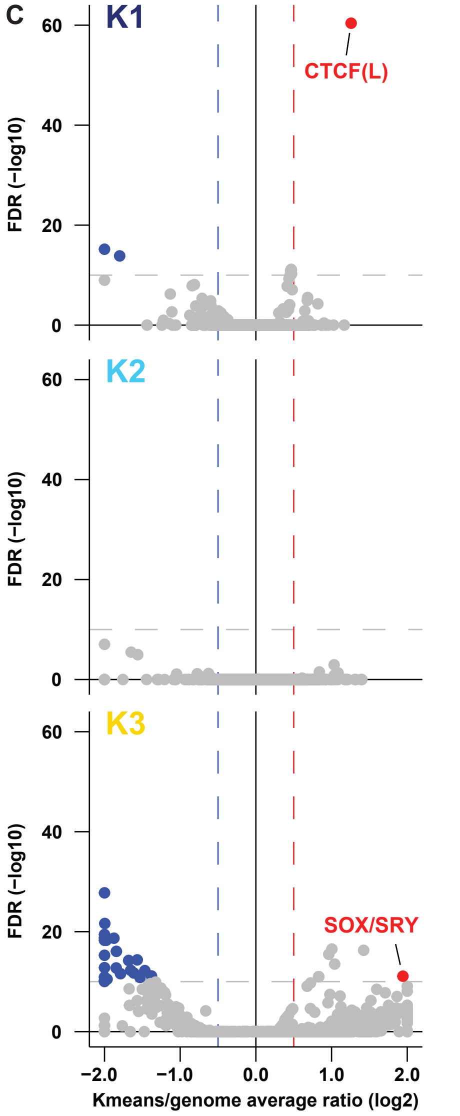


**D** Top 5 most significant biological processes (3 Kmeans clusters)

| ID | Biological process | Fold change | FDR |
|---|---|---|---|
| **K1 specific GO term:** | | | |
| GO:0043271 | negative regulation of ion transport | 2.78 | 2.72E-02 |
| **K2 specific GO term:** | | | |
| GO:0071824 | protein-DNA complex subunit organization | 4.76 | 6.16E-08 |
| GO:0065004 | protein-DNA complex assembly | 5.04 | 2.97E-07 |
| GO:0034728 | nucleosome organization | 5.36 | 2.51E-06 |
| GO:0006334 | nucleosome assembly | 5.97 | 5.59E-06 |
| GO:0031497 | chromatin assembly | 5.25 | 5.59E-06 |
| **K3 specific GO term:** | | | |
| GO:0051056 | regulation of small GTPase mediated signal transduction | 3.88 | 2.82E-13 |
| GO:0051241 | negative regulation of multicellular organismal process | 2.43 | 1.84E-12 |
| GO:0051093 | negative regulation of developmental process | 2.43 | 7.51E-11 |
| GO:0046578 | regulation of Ras protein signal transduction | 4.04 | 4.01E-10 |
| GO:0022603 | regulation of anatomical structure morphogenesis | 2.29 | 8.06E-10 |
